# Effect of high-altitude exposure on skeletal muscle mitochondrial subcellular distribution, ultrastructure and respiration in sea-level residents

**DOI:** 10.1101/2024.10.11.617733

**Authors:** Camilla Tvede Schytz, Joachim Nielsen, Niels Ørtenblad, Anne-Kristine Meinild Lundby, Robert Acton Jacobs, Carsten Lundby

## Abstract

Endurance exercise performance is associated with a well-developed skeletal muscle mitochondrial network, which is composed of subsarcolemmal mitochondria interconnected with intermyofibrillar mitochondria bending around the myofibrils. High-altitude exposure is typically incorporated in elite sport training regimens, but little is known about how this network adapts to an environment characterised by tissue hypoxia. For this reason, we investigated how high-altitude exposure affects mitochondrial subcellular distribution, ultrastructure, respiratory control, and intrinsic mitochondrial respiratory capacity. Nine healthy and recreationally active sea-level residents (eight males and one female) resided at an altitude of 3454 m with biopsies collected from the vastus lateralis muscle before and after 7 and 28 days at high altitude. The muscular mitochondrial volume density (*MitoVD*) increased after high-altitude exposure, driven by an increase in the intermyofibrillar *MitoVD*. This was however accompanied by a decreased cristae surface area per skeletal muscle fibre volume (*MuscularCD*) because of a decline in the cristae surface area per mitochondrial volume (*MitoCD*). Despite a reduced *MuscularCD*, mass-specific maximal coupled respiration (*OXPHOS_CII+CI+ETF*) increased slightly, and was considerably elevated when normalised to *MuscularCD*, suggesting intrinsic adaptations to high altitude. The difference between cristae-specific *OXPHOS_CII+CI+ETF* and an associated cristae-specific leak respiration (*Leak_CII+CI+ETF*) indicated a markedly higher degree of coupling between the electron flow in the electron transport system and ATP production. As the effect size 95% confidence intervals includes trivial effects the results need to be substantiated. In conclusion, high-altitude exposure altered mitochondrial subcellular distribution, ultrastructure and induced intrinsic respiratory adaptations.

## Introduction

Skeletal muscle mitochondria are important for endurance exercise performance in athletes (Jacobs *et al*., 2011) and are most often classified in two subcellular localisations: one surrounding the myofibrils in the intermyofibrillar space and another localised in the subsarcolemmal space between sarcolemma and the outermost myofibrils (Eisenberg, 1983). Here, the intermyofibrillar pool is the significantly largest making up around 80-85% of the total pool (Hoppeler *et al*., 1985; Rösler *et al*., 1985; Schytz *et al*., 2024). While endurance athletes may often make use of high-altitude training camps, knowledge about how high-altitude environments, characterised by tissue hypoxia, affects the mitochondrial network and its subcellular distribution is limited.

Only few studies have investigated how high-altitude exposure affects skeletal muscle mitochondrial volume density, which is the mitochondrial volume per fibre volume (muscular *MitoVD*) and how this manifest on the subcellular level. Exposing Caucasians to high altitudes for ∼5-8 weeks during climbing expeditions has been shown to decrease muscular *MitoVD* by approximately 20% with a larger relative drop in the subsarcolemmal than the intermyofibrillar *MitoVD* (Hoppeler *et al*., 1990; Levett *et al*., 2012). On the contrary, no change was observed in the intermyofibrillar *MitoVD* after a 40-day residence in a hypobaric chamber where pressure was gradually adjusted toward Mt. Everest equivalent altitude (MacDougall *et al*., 1991). However, field expeditions and pressure chamber residence are exposed to factors such as inconsistent caloric intake and exercise patterns different from habitual activity, which may confound these results. Taking these issues into account, we have previously demonstrated that exposing sea-level residents to 3454 m altitude for 28 days induced a small increase in muscular *MitoVD* due to an augmentation of the intermyofibrillar pool (Jacobs *et al*., 2016). Since a high-altitude environment is characterised by tissue hypoxia, high-altitude exposure may also alter mitochondrial respiratory efficiency.

Mitochondrial respiratory control is evaluated using high-resolution respirometry. Studies show trivial effects on maximal coupled respiration normalised to overall tissue sample mass (i.e. mass- specific) when Caucasians are exposed to high altitude for 16-59 days (Jacobs *et al*., 2013; Horscroft *et al*., 2017; Chicco *et al*., 2018). To evaluate intrinsic mitochondrial respiratory capacity, these mass- specific respiratory measures can be combined with quantitative measures defining the skeletal muscle mitochondrial content. Here, muscular *MitoVD* has typically been applied or commonly used biomarkers hereof such as citrate synthase (CS) activity. Accordingly, Jacobs *et al*. (2012) found a ∼30% decrease in maximal coupled respiration normalised to CS activity after a 28-day sojourn at 3454 m. However, assessing muscular *MitoVD* alone may not fully capture the skeletal muscle mitochondrial content when evaluating its intrinsic respiratory capabilities (Nielsen *et al*., 2017; Schytz *et al*., 2024).

Structurally, mitochondria consist of both an outer and an inner membrane where the inner membrane is characterised by infoldings known as cristae (Palade, 1952, 1953; Sjöstrand, 1953). Most of the complexes responsible for energy transduction are localised in the cristae membrane (Gilkerson *et al*., 2003; Schlame, 2021). Consequently, the ATP-producing capability of mitochondria should inherently be dependent on the cristae surface area per mitochondrial volume referred to as mitochondrial cristae density (*MitoCD*). As a result, respiration per cristae surface area may provide a more accurate evaluation of intrinsic mitochondrial respiratory capacity than respiration per mitochondrial volume. Since traditional high-resolution respirometry are performed on skeletal muscle samples and not isolated mitochondria the cristae surface area has to be expressed specific to the skeletal muscle volume rather than the mitochondrial volume. Thus, the muscular cristae density, which is the cristae surface area per fibre volume (*MuscularCD*), may serve as a more complete quantitative structural measure of skeletal muscle mitochondrial content than muscular *MitoVD* alone. In support of this, we recently demonstrated that differences in mass-specific maximal coupled respiration across individuals of a broad range of training statuses were attributed to variations in *MuscularCD* and not intrinsic adaptations (Schytz *et al*., 2024). However, no study has investigated the effect of high-altitude exposure on *MuscularCD* and therefore knowledge on how high altitude affects mitochondrial respiratory efficiency is very limited. Therefore, based on human skeletal muscle biopsies from Jacobs *et al*. (2016), the aim of this study was to investigate the effect of high- altitude exposure on mitochondrial subcellular distribution (*MitoVD*s), ultrastructure (*MitoCD* and *MuscularCD*), respiratory control and intrinsic mitochondrial respiratory capacity by combining mass-specific mitochondrial respiration with *MuscularCD*.

## Methods

### Ethical approval

The human skeletal muscle biopsy material included in the present study have in part been published previously (Jacobs *et al*., 2016; Schytz *et al*., 2024). This study was approved by the ethical committee of the Eidgenössische Technische Hochschule in Zürich (ETH, EK 2011-N-51) and conformed to the standards set by the *Declaration of Helsinki*. Participants were informed about the experiments and potential risks before providing their written informed content.

### Participants, study overview and skeletal muscle biopsies

Nine healthy and recreationally active sea-level residents (eight males and one female) were enrolled (median (25th-75th percentile): age, 27 years (23-29); body mass (BM), 76 kg (73-81); height, 1.81 m (1.78-1.83), and maximal oxygen consumption (V̇O_2max_), 50.5 (48.6-58.9) mL O_2_ x min^-1^ x kg^-1^ BM).

In order to study the effect of high altitude on skeletal muscle mitochondrial subcellular distribution, ultrastructure and respiration, the participants were exposed to high altitude in the Swiss Alps for 28 days (Fig. 1A). Participants replicated normal sea level activities to the best of their ability at the research station where a Monark ergometer (Monark, E839, Varberg, Sweden) and six private dumbbells were brought to be able to mimic daily commuting (*n*=2) and two sessions (30-35 min) of upper body strength training per week (*n*=1) (Jacobs *et al*., 2016).

**Figure 1:**
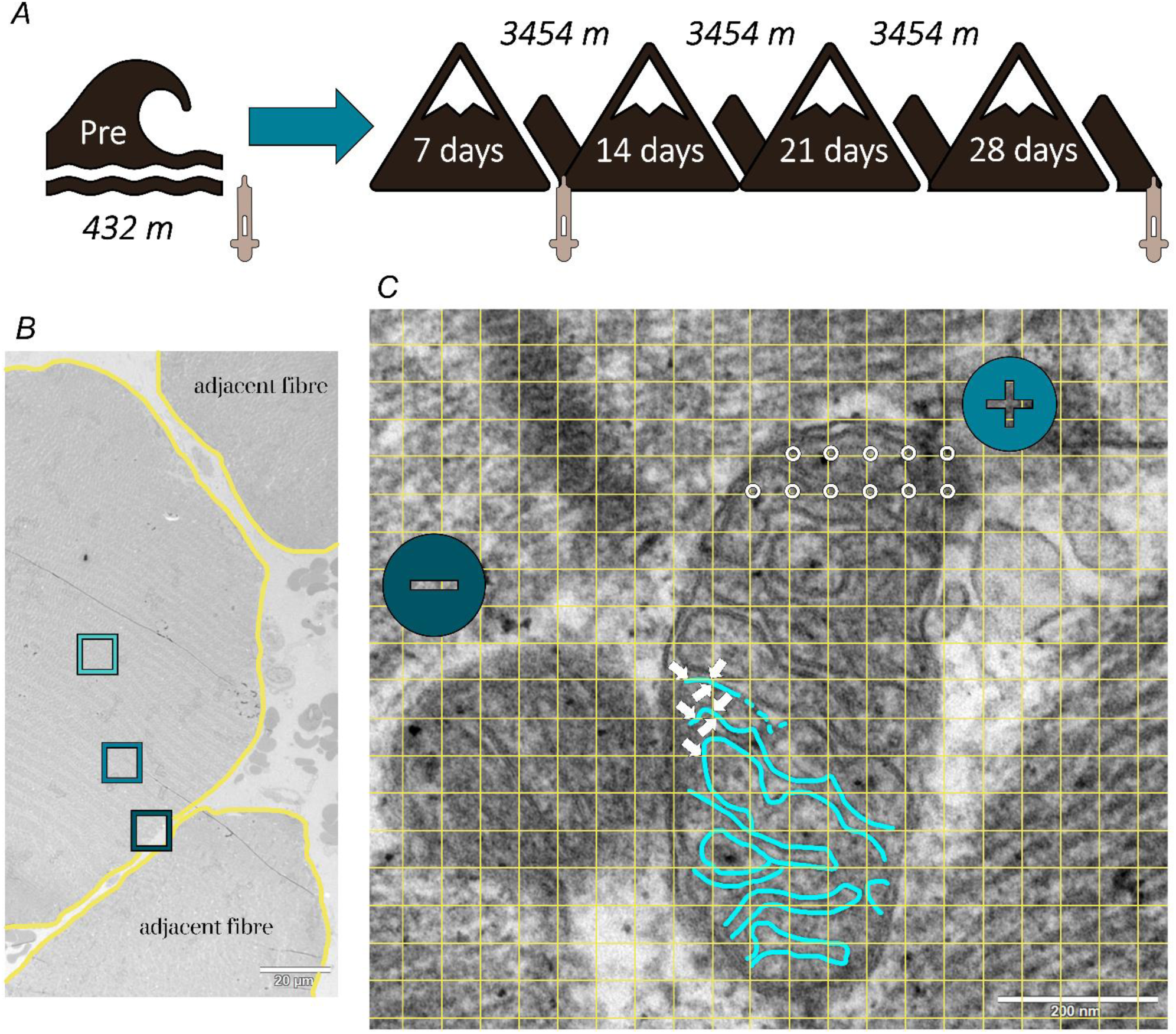
Study design and estimation of mitochondrial cristae density. *A*, skeletal muscle biopsies were obtained at sea level (432 m) (Pre) and following 7 and 28 of high-altitude (3454 m) exposure. *B*, at least three images (boxes) were obtained along the depth of each skeletal muscle fibre meaning from just beneath the sarcolemma (yellow line) to the superficial and central parts of the myofibrillar space. *C*, each image included at least one mitochondrial profile of acceptable quality (plus icon) defined as clear visibility and with no or few missing traces of cristae (part of the trace highlighted in blue), while the remaining profiles (minus icon) were discarded. Grids (yellow) were overlayed each image. Number of cristae intersections with horizontal and vertical lines (arrows) in a grid of known grid line distance were used to estimate the cristae surface area, while number of hits of the mitochondria (circles) in a grid of a known set of points gives the mitochondrial volume. Then, the ratio between the cristae surface area and the mitochondrial volume gives the *MitoCD*.

Skeletal muscle biopsies were obtained from the *m. vastus lateralis* at sea level in Zürich (432 m) prior to the high-altitude ascent (Pre) and following 7 and 28 days of exposure to high altitude at the Jungfraujoch Research Station (3454 m) (Fig. 1A). Biopsies were obtained at rest using the Bergström technique with suction under local anaesthesia (1% lidocaine) of the skin and superficial muscle fascia. The skeletal muscle biopsies were dissected free of fat and connective tissue and divided into specimens used in this study for analysis by transmission electron microscopy (TEM) and high- resolution respirometry. Part of data have already been presented in accompanying papers: V̇O_2max_, *MitoVD* (muscular, intermyofibrillar and subsarcolemmal) and mitochondrial respiration (but not normalised values) in Jacobs *et al*. (2016) and pre-altitude *MitoCD* and *MuscularCD* in Schytz *et al*. (2024). The de novo analysis in this study is high-altitude *MitoCD*, *MuscularCD* and normalised respirations values throughout acclimatisation. Furthermore, the analysis is based on effect sizes and associated 95% confidence interval contrary to previous analysis.

### Maximal oxygen consumption

V̇O_2max_ was determined in an incremental cycling test by indirect calorimetry as previously described in Jacobs *et al*. (2016), where V̇O_2max_ was determined as the highest mean across a 10 s period.

### MitoVD, MitoCD and MuscularCD

#### Preparation of fixed muscle fibres for transmission electron microscopy

Four randomly chosen specimens (∼1x1x1 mm in size) from each skeletal muscle biopsy were prepared for TEM as previously described in Schytz *et al*. (2024). Thus, the specimens were fixed with 2.5% glutaraldehyde in 0.1 M sodium cacodylate buffer (pH 7.3), stored at room temperature for 24 h, then washed three times in 0.1 M sodium cacodylate buffer and stored in 0.1 M sodium cacodylate buffer at 4 °C until further processing. These fixed muscle specimens were post-fixated with 1% osmium tetroxide in 0.1 M cacodylate buffer for 120 min at room temperature. Subsequently, the muscle specimens were washed three times with milliQ H_2_O followed by a block-staining with 2% uranyl acetate in milliQ H_2_O overnight at room temperature. Afterwards graded dehydration was accomplished in a tissue processor (Leica EM TP, Leica Microsystems, Germany): 10 min 70% ethanol, 15 min 96% ethanol, and 4x30 min 100% ethanol. Then the muscle specimens were infiltrated with graded mixtures of propylene oxide and Epon (2x30 min propylene oxide, 60 min 1:1 propylene oxide:Epon, overnight 1:1 propylene oxide:Epon, 180 min 100% Epon). Next, each muscle specimen was embedded in 100% Epon in an individual block in a non-oriented (isotropic) manner and then Epon was polymerised for 48 h at 60 °C. Finally, 3-4 ultra-thin (∼65 nm) sections were cut in two depths all separated by 20 μm from each block and placed on two individual 50 mesh hexagonal copper grids (Plano GmbH, Wetzlar, Germany) before post-staining the sections with Reynolds lead citrate.

#### Image capturing by transmission electron microscopy

To determine *MitoVD*, the ultra-thin contrasted sections were photographed in a random and systematic order using a FEI Tecnai G2 Spirit transmission electron microscope (FEI, Hillsboro, OR, USA) mounted with an Orius SC1000 CCD camera (Gatan, Pleasanton, CA, USA). From each biopsy, 216 images (3840x2528 pixels, with one pixel being 4.14x4.14 nm) were obtained distributed on 24 meshes (9 images per mesh) from 8 grids originating from 4 blocks (i.e. specimens). The images were acquired using the automated image capturing function in the TEM photomontage software by selecting a random start point for the first image, and taking the subsequent images at a fixed x-,y- intervals of – 400%.

To determine *MitoCD*, the ultra-thin contrasted sections were re-photographed at a greater magnification (46000×) using a pre-calibrated Philips CM100 transmission electron microscope (Philips, Eindhoven, The Netherlands) and an Olympus Veleta camera (Olympus Soft Imaging Solutions, Münster, Germany). Two blinded investigators photographed 8-17 fibres per biopsy including at least one image from the central, superficial and subsarcolemmal region of the fibre (Fig. 1B). These images contained a minimum of one mitochondrial profile of acceptable quality, meaning a clear visibility and with no or few missing traces of the inner mitochondrial membrane (Fig. 1C).

*MitoVD*, *MitoCD* and the combined measure, *MuscularCD*, were then determined using stereology, which is a body of mathematical models enabling quantifying 3D objects of a given structure based on measurements carried out on 2D images on sections of this structure (Weibel, 1980).

#### Stereological analysis to assess MitoVD

Mitochondrial volume density was estimated by point counting as stereological principles dictate that *V_V_*=*A_A_*=*P_P_*, permitting that a volume density (*V_V_*) can be estimated from a point density (*P_P_*) where *A_A_* is an area density (see proofs in Weibel (1980)). This was accomplished by overlaying grids with a spacing of 1x1 µm onto each image using the Cavalieri feature in the Stereo-Investigator software (MBF Bioscience, USA). Each point was defined as either intermyofibrillar mitochondria, subsarcolemmal mitochondria, or skeletal muscle, and points not belonging to the muscle specimen, due to the random image acquisition, were excluded. Mitochondria boundaries were identified at 8200× magnification. Intermyofibrillar and subsarcolemmal *MitoVD* were determined, and the sum make up the mitochondrial volume in total per fibre volume (muscular *MitoVD*).

#### Stereological analysis to assess MitoCD and MuscularCD

*MitoCD* (*S_V_*) was estimated by point counting and counting intersections with lines for each mitochondrial profile (Fig. 1C) as previously described in Schytz *et al*. (2024). This was accomplished using the stereological formular: *S_V_* = 2 x *I_dbl_* x (*P_mi_* x *k* x *d*)^-1^ where *I_dbl_* is the number of times the inner membrane intersects the lines of the test system, *P_mi_* is the number of points in the test system that hits the mitochondrial profile, while the constants *k* and *d*, are characteristics of the test system (see mathematical derivation of this formular in Weibel (1980)). Grid sizes of 270 x 270 nm and 90 x 90 nm were applied to measure *I_dbl_* and *P_mi_*, respectively. The mitochondrial profiles were analysed by one blinded investigator in a randomised order.

Since mitochondrial profiles had to be of acceptable quality to be included in the analysis, only parts of the profile were analysed for some mitochondrial profiles, while others were not analysed (Fig. 1C). In Schytz *et al*. (2024) an estimation of the coefficient of error (_est_CE) suggest that at least 8 profiles (whole or parts) must be included in the analysis to achieve a satisfactory high precision (_est_CE=0.1) of the *MitoCD*. Consequently, only biopsies with at least 8 analysed profiles were included, leading to an exclusion of two Pre-biopsies. Thus, sample size is lower for measures including *MitoCD*. The median number of mitochondrial profiles (whole or parts) analysed per participant was 33 (interquartile range: 24:41), and for the subcellular locations it was 14 (11:16), 12 (11:14) and 13 (11:15) for the central, superficial and subsarcolemmal region, respectively. All analyses were performed in ImageJ (ImageJ 1.53e, National Institutes of Health, USA).

The cristae surface area per fibre volume (*MuscularCD*) were determined as the product between muscular *MitoVD* and *MitoCD*.

### Mitochondrial respiration

#### Preparation of permeabilised muscle fibres for high-resolution respirometry

Muscle specimens were prepared as described in Jacobs *et al*. (2012). A ∼20 mg specimen from each skeletal muscle biopsy was placed in an ice-cold biopsy preservation solution (BIOPS: 2.77 mM CaK_2_EGTA buffer, 7.23 mM K_2_EGTA buffer, 0.1 µM free calcium, 20 mM imidazole, 20 mM taurine, 50 mM 2-(*N*-morpholino) ethanesulfonic acid hydrate (K-MES), 0.5 mM dithiothreitol (DTT), 6.56 mM MgCl_2_·6 H_2_O, 5.77 mM ATP, and 15 mM phosphocreatine; pH: 7.1) immediately after biopsy extraction. The preparation was initiated by a gentle mechanical dissection of the skeletal muscle specimens with the tip of two 18-gauge needles to achieve a high degree of fibre separation, which was verified microscopically. Then chemically permeabilisation of the skeletal muscle fibres was accomplished by incubating the specimens in 2 mL BIOPS with saponin (50 µg·mL^-1^) for 30 min at 4 °C. Subsequently, the skeletal muscle specimens were washed for 10 min at 4 °C in mitochondrial respiration medium 05 (MiR05: 0.5 EGTA, 3 mM MgCl_2_ · 6 H_2_O, 60 mM potassium lactobionate, 20 mM taurine, 10 mM KH_2_PO_4_, 20 mM HEPES, 110 mM sucrose, and 1 mg·mL^-1^bovine serum albumin (BSA); pH: 7.1). Next, mass measurements were conducted by first blotting ∼1-3 mg bundles of fibres carefully on a filter paper by placing them for 5 s onto the paper with a sharp pair of forceps (angular tip), and in the meantime, liquid was removed from the tip of the forceps with a dry filter paper. Next, the fibre bundles were lifted from the filter paper and while holding them with the forceps the bundles were briefly touched once more onto a dry area of the filter paper. Then the fibre bundles were immediately placed onto a tarred balance-controlled scale (XS205, DualRange Analytical Balance; Mettler-Toledo AG, Greifensee, Switzerland). Finally, after determining the wet weight (ww) the fibre bundles were immediately transferred into the Oxygraph- 2k chamber.

#### High-resolution respirometry

Mitochondrial respiration was analysed by measuring oxygen consumption when adding substrates, uncouplers and inhibitors to the chambers of a high-resolution Oxygraph-2k (Oroboros, Innsbruck, Austria). This *Substrate*-*Uncoupler*-*Inhibitor Titration* (SUIT) protocol (Gnaiger, 2014) were conducted with titrations in series in the following order: **(1)** malate (2 mM) and octanoyl carnitine (0.2 mM) to assess leak respiration (*Leak_ETF+CI*) (state 2) supported by an electron flow through Complex I and a transfer of electrons from the electron-transferring flavoprotein (ETF) to ETF:ubiquinone oxidoreductase that conveys electrons to the ubiquinone pool of the electron transport system (ETS) which in turn relays electrons to Complex III; **(2)** saturating ADP (5 mM) to couple the electron flow to the phosphorylation of ADP to ATP to assess ETF Complex I-linked coupled respiration as a measure of fatty acid oxidative capacity (*FAO_ETF+CI*) (state 3). In this state, titration of malate (added in the previous step) is necessary to prevent blocking the β-oxidation by an accumulation of acetyl CoA; **(3)** lactate (30 mM), pyruvate (5 mM) and glutamate (10 mM) to achieve maximal electron flow through Complex I and succinate (10 mM) to provide substrate for Complex II to assess Complex II-I-ETF-linked oxidative phosphorylation capacity (*OXPHOS_CII+CI+ETF*) (state 3); **(4)** cytochrome c (10 µM) to assess the integrity of the outer mitochondrial membrane (Kuznetsov *et al*., 2008) where a >20% increase in respiration was considered as a sign of damage leading to exclusion of data; **(5)** oligomycin (2.5 µM) to inhibit the ATP synthase (Complex V) to assess Complex II-I-ETF-supported leak respiration (*Leak_CII+CI+ETF*) (state 4o); **(6)** titration of the proton ionophore carbonyl cyanide *p*- (trifluoromethoxy) phenylhydrazone (FCCP, 1-2.5 µM) to uncouple the ATP synthase (Complex V) from the ETS; **(7)** rotenone (0.5 µM) to inhibit Complex I and antimycin A (2.5 µM) to inhibit Complex III to block electron flow to assess residual oxygen consumption (ROX), indicative of non- mitochondrial oxygen consumption. All measurements were corrected for ROX. To examine potential intrinsic adaptations, we assessed the coupling efficiency between the suggested electron flow in the ETS and ATP production via oxygen kinetics by subtracting mass-, mitochondrial- and cristae-specific leak respiration from mass-, mitochondrial-, and cristae-specific coupled respiration, respectively. Leak and coupled respiration were supported by identical substrates. This measure was preferred since this, as opposed to the typical calculated respiratory control ratio, is dependent on the normalising procedure, which is of importance as this affects the interpretation of intrinsic adaptations (Schytz *et al*., 2024).

Mitochondrial respiration measurements were performed in MiR05 plus catalase (280 IU x mL^-1^). All measurements were analysed in duplicates at 37 °C and conducted in a hyper-oxygenated environment within the chambers (200 to 450 nmol O_2_ x mL^-1^) to avoid potential oxygen diffusion limitations. Standardised instrumental calibrations were conducted to correct for oxygen consumption by the sensor and chemical medium, exterior leak and diffusion of oxygen back into the chamber from various components (Jacobs *et al*., 2012). Rate of respiration was resolved by software (DatLab, Oxygraph 2k, Oroboros, Innsbruck, Austria), accounting for non-linear alterations in the negative time derivative of the rate of oxygen respiration (Jacobs *et al*., 2012). Rate of respiration is expressed in pmol O_2_ x s^-1^x mg ww^-1^.

### Statistics

The effect of high altitude on the parameters was investigated by estimating the distribution of the bootstrapped-resampled paired mean difference between both 7 and 28 days of high-altitude exposure and the Pre high-altitude time point (control) using estimationstats.com (Ho *et al*., 2019). 5000 bootstrap samples were taken; the 95% confidence interval is bias-corrected and accelerated.

## Results

As reported previously (Jacobs *et al*., 2016), the effect of high altitude on muscular *MitoVD* was dependent on both the duration of the high altitude and on the subcellular location within the skeletal muscle fibres (Fig. 2). While the *MitoVD* within the intermyofibrillar space increased (10%) after 28 days at high altitude (Fig. 2B), the *MitoVD* in the subsarcolemmal space decreased after both 7 and 28 days (Fig. 2C). The wide 95% confidence interval of the effect size of the latter, however, suggest an uncertain result. Since the muscular *MitoVD* is largely dependent on the *MitoVD* of intermyofibrillar mitochondria, it also increased (6%) after 28 days at high altitude (Fig. 2A).

**Figure 2:**
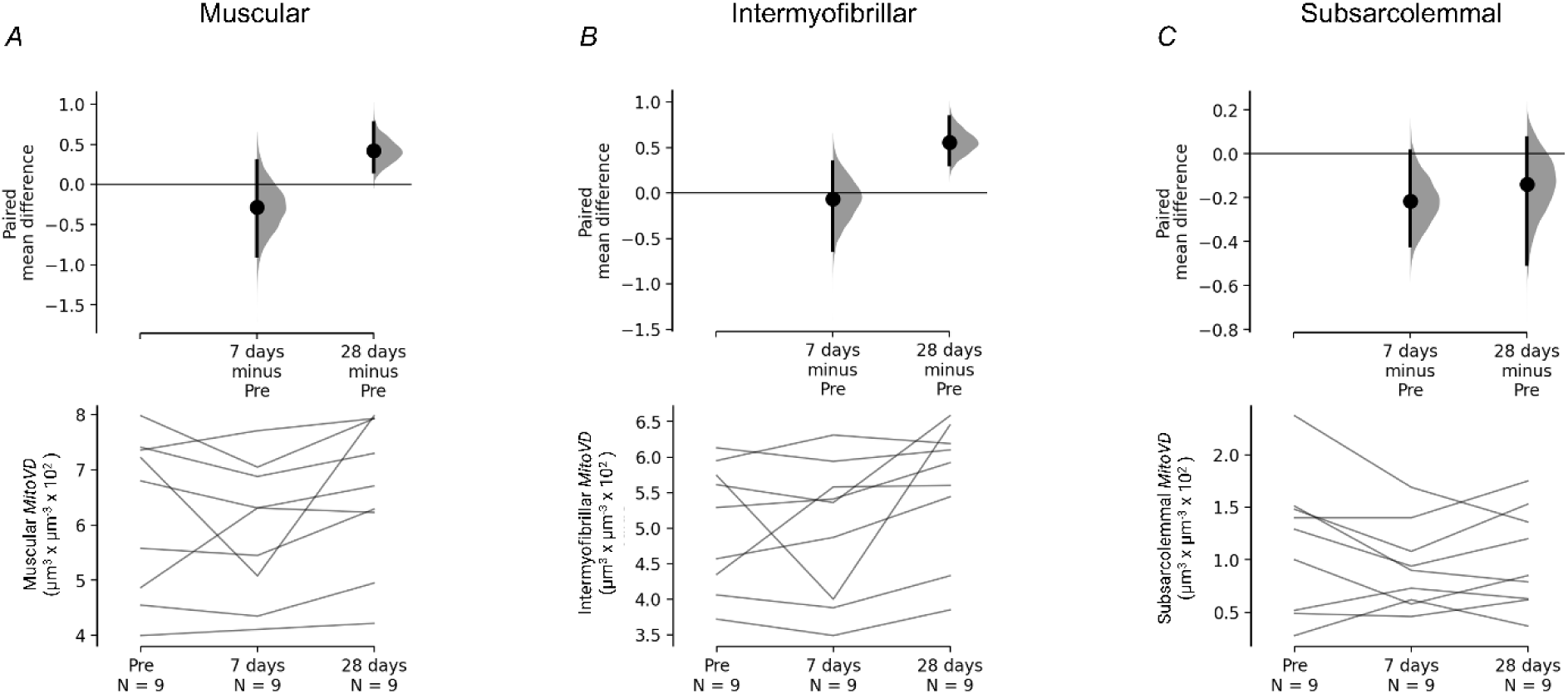
Effect of high altitude on muscular and subcellular MitoVD. *A*, muscular *MitoVD*; *B*, intermyofibrillar *MitoVD*; *C*, subsarcolemmal *MitoVD*. The paired mean difference between the timepoint before high-altitude exposure (Pre) and after high-altitude exposure (7 and 28 days) is displayed in a Cumming estimation plot. The top panels show the distribution of the bootstrapped-resampled paired mean difference between both the 7 days and 28 days timepoints and the Pre high-altitude timepoint. In each effect size half-violin plot, the black dot denotes the mean of the distribution, and the black vertical bars indicate 95% confidence intervals. The bottom panels consist of a slopegraph whereby each line represents a set of observations from an individual.

*MitoCD* decreased (5-10%) after 28 days at high altitude (Fig. 3A). Thus, despite an increased muscular *MitoVD*, the *MuscularCD* did not increase with acclimatiation to high altitude. The effect sizes suggest a small decrease (5-10%) in *MuscularCD* following 7 and 28 days at high altitude, but a large decrease is also compatible with the 95% confidence intervals, especially after 7 days at high altitude (Fig. 3B).

**Figure 3:**
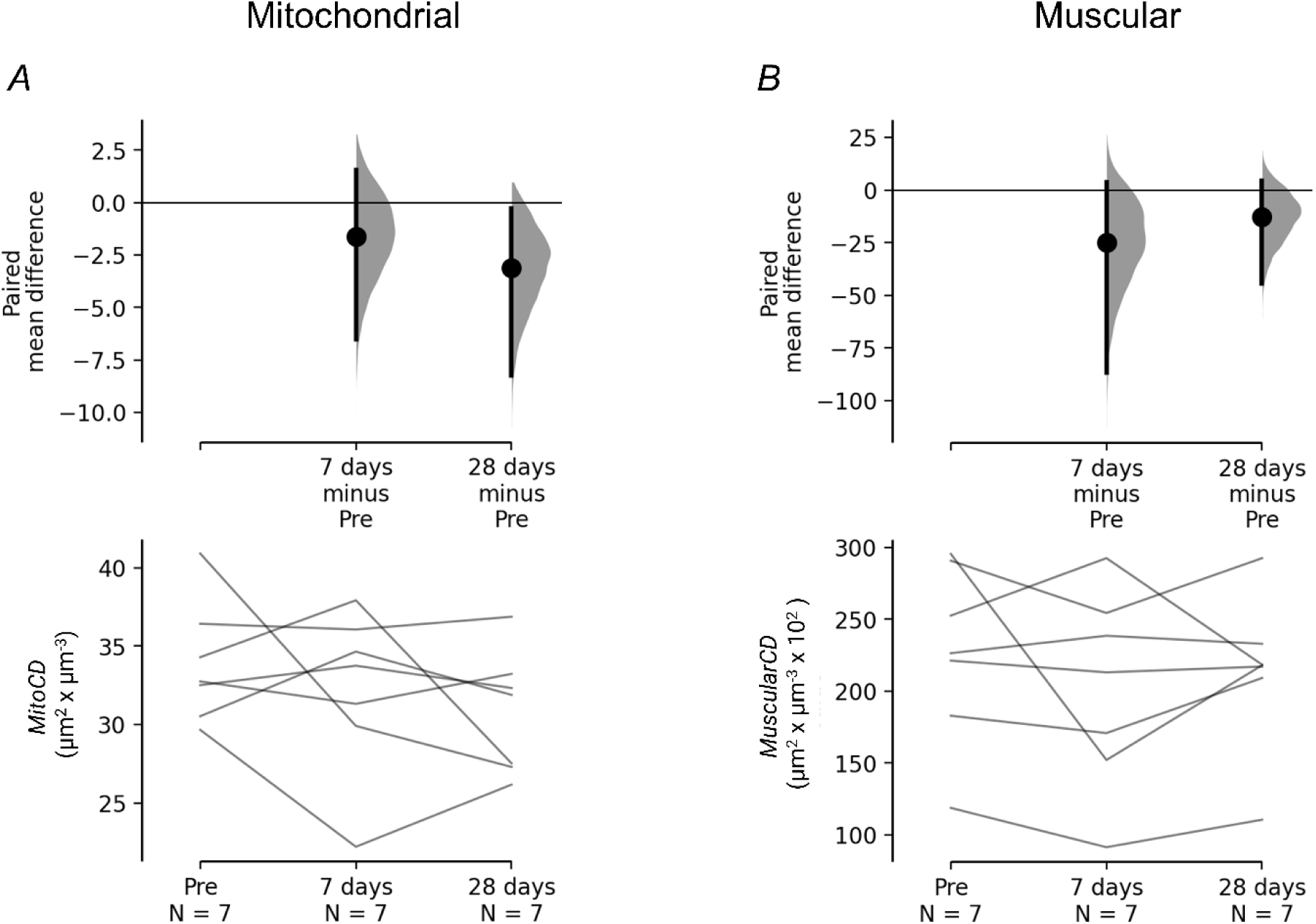
Effect of high altitude on MitoCD and MuscularCD. *A, MitoCD*; *B*, *MuscularCD*. The paired mean difference between the timepoint before high-altitude exposure (Pre) and after high-altitude exposure (7 and 28 days) is displayed in a Cumming estimation plot. The top panels show the distribution of the bootstrapped-resampled paired mean difference between both the 7 days and 28 days timepoints and the Pre high-altitude timepoint. In each effect size half-violin plot, the black dot denotes the mean of the distribution, and the black vertical bars indicate 95% confidence intervals. The bottom panels consist of a slopegraph whereby each line represents a set of observations from an individual.

The indicated decrease in *MuscularCD* affects the interpretation of the mitochondrial respiration data. Thus, when *OXPHOS_CII+CI+ETF* and *Leak_CII+CI+ETF* are normalised to the yielded muscular wet weight and to the muscular *MitoVD* only small increases (5-15%) are suggested by the effect sizes and corresponding 95% confidence intervals after 28 days at high altitude (Fig. 4A-B, D-E). However, when normalising to *MuscularCD*, *OXPHOS_CII+CI+ETF* increased by 25% after 7 and 28 days at high altitude where increases of 5-60% are compatible with the 95% confidence intervals (Fig. 4C). This increase was accompanied by skewed effect size 95% confidence intervals towards an increased cristae-specific *Leak_CII+CI+ETF* after 7 (10%) and 28 days (20%) at high altitude (Fig. 4F). To investigate the coupling degree between the suggested electron flow and the ATP production via oxygen kinetics, cristae-specific *Leak_CII+CI+ETF* was subtracted from cristae- specific *OXPHOS_CII+CI+ETF*. Despite an increased leak, a 30-40% higher degree of mitochondrial coupling per *MuscularCD* was indicated following 7 and 28 days at high altitude, which was not evident in the mass- and mitochondrial-specific data (Fig. 5).

**Figure 4:**
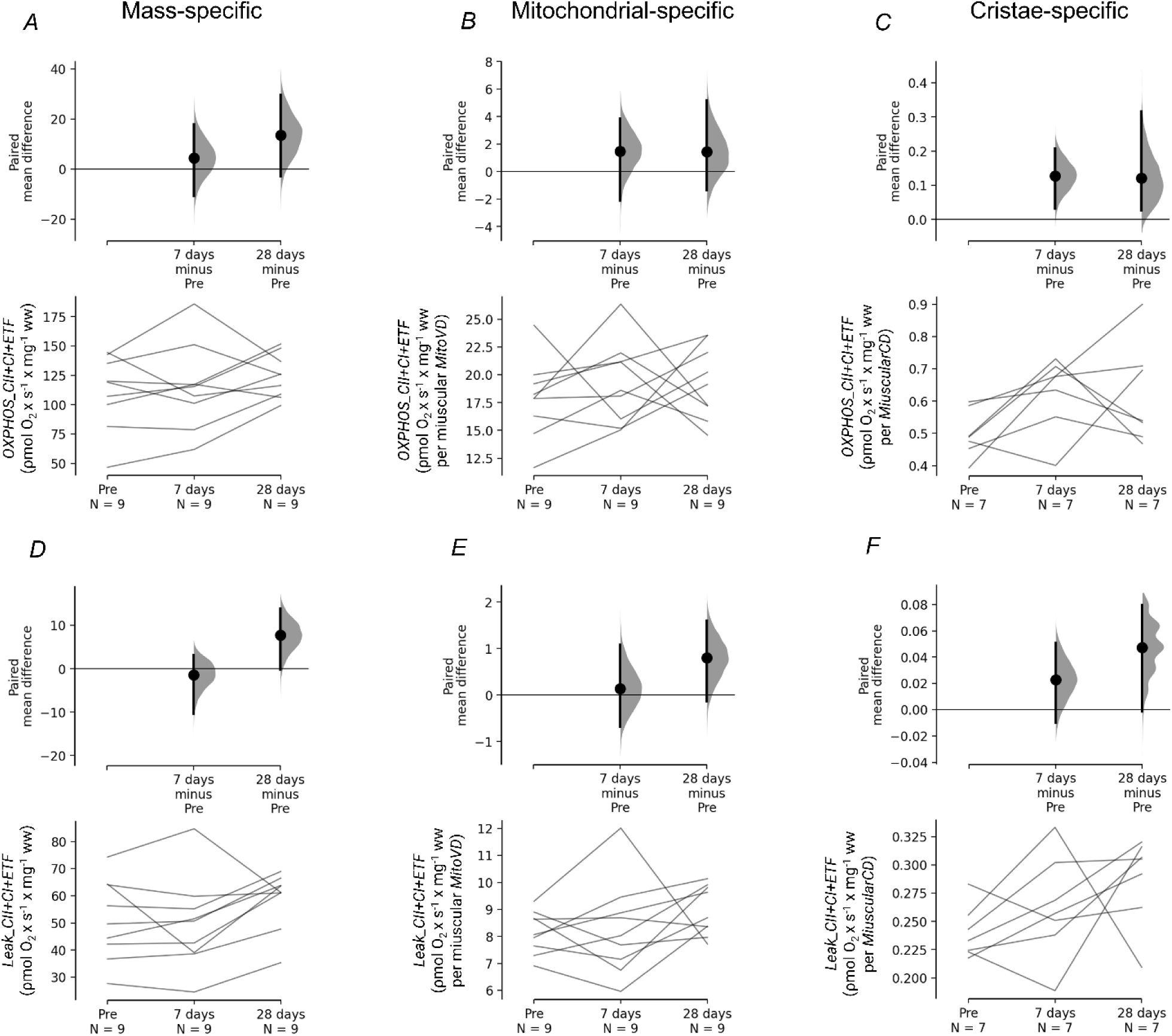
Effect of high altitude on mass-, mitochondrial- and cristae-specific *OXPHOS_CII+CI+ETF* and *LEAK_CII+CI+ETF*. *A,* mass-specific *OXPHOS_CII+CI+ETF* (respiration normalised to the wet weight of the muscle specimen); *B*, mitochondrial-specific *OXPHOS_CII+CI+ETF* (mass-specific respiration normalised to muscular *MitoVD*); *C*, cristae-specific *OXPHOS_CII+CI+ETF* (mass-specific respiration normalised to *MuscularCD*). *D,* mass-specific *LEAK_CII+CI+ETF*; *E*, mitochondrial-specific *LEAK_CII+CI+ETF*; *F*, cristae-specific *LEAK_CII+CI+ETF*. The paired mean difference between the timepoint before high-altitude exposure (Pre) and after high-altitude exposure (7 and 28 days) is displayed in a Cumming estimation plot. The top panels show the distribution of the bootstrapped-resampled paired mean difference between both the 7 days and 28 days timepoints and the Pre high-altitude timepoint. In each effect size half-violin plot, the black dot denotes the mean of the distribution, and the black vertical bars indicate 95% confidence intervals. The bottom panels consist of a slopegraph whereby each line represents a set of observations from an individual.

**Figure 5:**
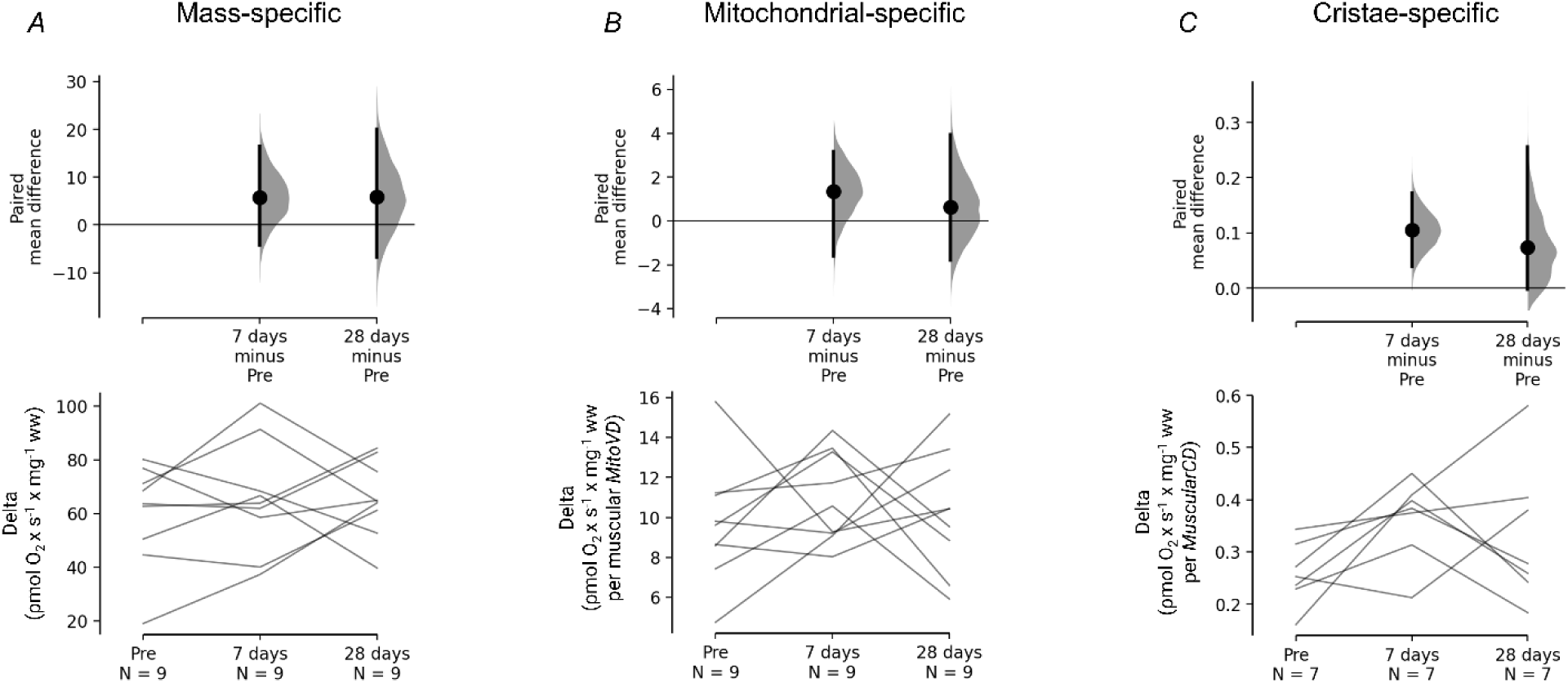
Effect of high altitude on mass-, mitochondrial- and cristae-specific mitochondrial coupling degree. *A,* mass-specific *OXPHOS_CII+CI+ETF* minus mass-specific *LEAK_CII+CI+ETF* (delta); *B,* mitochondrial-specific *OXPHOS_CII+CI+ETF* minus mitochondrial-specific *LEAK_CII+CI+ETF* (delta); *C,* cristae-specific *OXPHOS_CII+CI+ETF* minus cristae-specific *LEAK_CII+CI+ETF* (delta). The paired mean difference between the timepoint before high-altitude exposure (Pre) and after high-altitude exposure (7 and 28 days) is displayed in a Cumming estimation plot. The top panels show the distribution of the bootstrapped-resampled paired mean difference between both the 7 days and 28 days timepoints and the Pre high-altitude timepoint. In each effect size half-violin plot, the black dot denotes the mean of the distribution, and the black vertical bars indicate 95% confidence intervals. The bottom panels consist of a slopegraph whereby each line represents a set of observations from an individual.

The effect of high altitude on mitochondrial fatty acid oxidation was dependent on the duration of the exposure. Thus, only trivial effects are seen for mass-, mitochondrial-, and cristae-specific *FAO_ETF+CI* after 7 days at high altitude, while the effect size 95% confidence intervals are skewed towards a 10-15% decrease after 28 days (Fig. 6A-C). Together with an augmented and unaltered *Leak_ETF+CI* after 7 and 28 days of high altitude, respectively, this led to a 15-25% decrease in mass- and mitochondrial-specific coupling degree (Fig. 6D-F and Fig. 7). However, this effect either vanished or was dampened when normalising to *MuscularCD* (Fig. 7).

**Figure 6:**
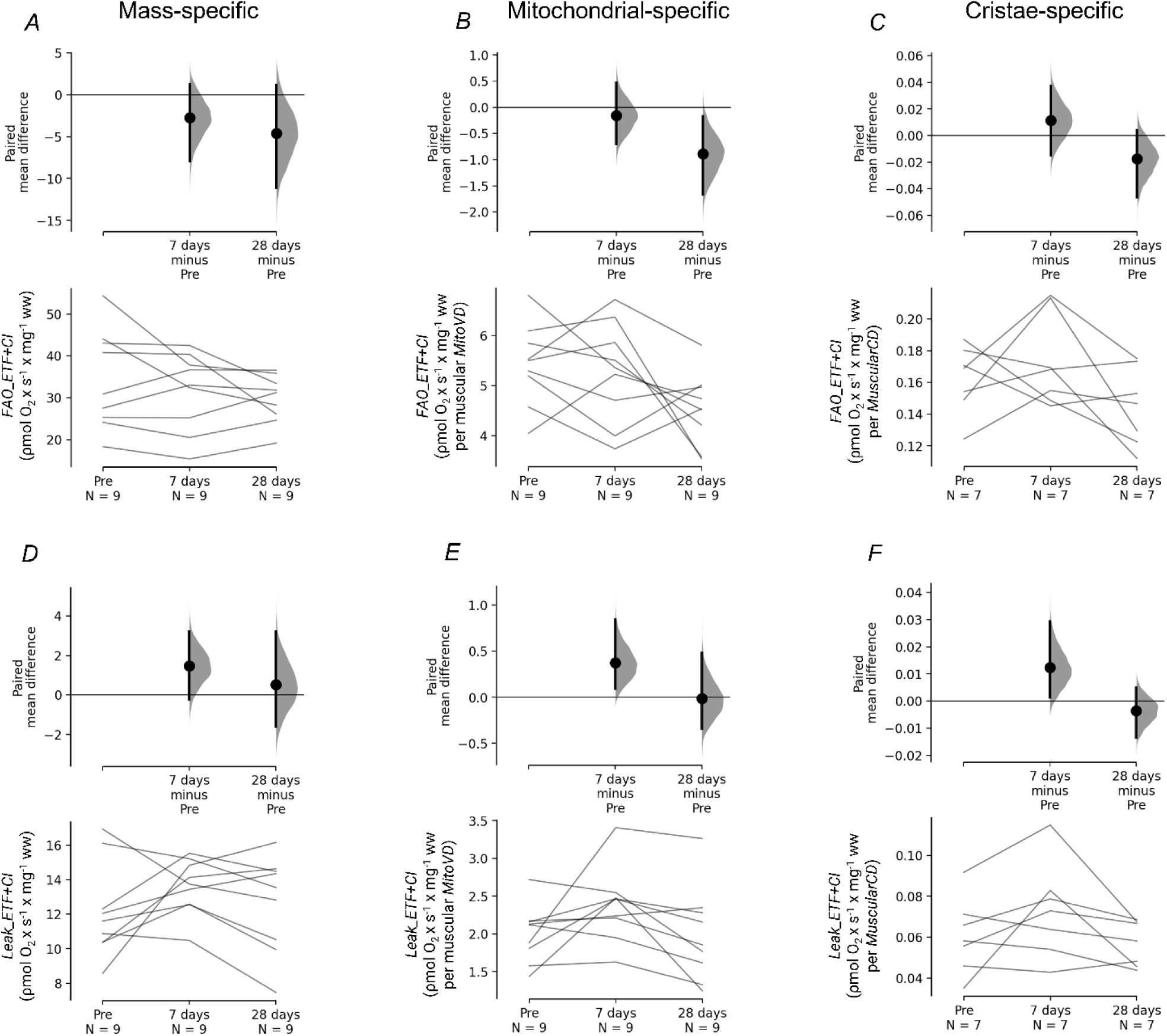
Effect of high altitude on mass-, mitochondrial- and cristae-specific *FAO_ETF+CI* and *LEAK_ETF+CI*. *A,* mass-specific *FAO_ETF+CI* (respiration normalised to the wet weight of the muscle specimen); *B*, mitochondrial-specific *FAO_ETF+CI* (mass-specific respiration normalised to muscular *MitoVD*); *C*, cristae-specific *FAO_ETF+CI* (mass-specific respiration normalised to *MuscularCD*). *D,* mass-specific *LEAK_ETF+CI*; *E*, mitochondrial-specific *LEAK_ETF+CI*; *F*, cristae-specific *LEAK_ETF+CI*. The paired mean difference between the timepoint before high-altitude exposure (Pre) and after high-altitude exposure (7 and 28 days) is displayed in a Cumming estimation plot. The top panels show the distribution of the bootstrapped-resampled paired mean difference between both the 7 days and 28 days timepoints and the Pre high-altitude timepoint. In each effect size half-violin plot, the black dot denotes the mean of the distribution, and the black vertical bars indicate 95% confidence intervals. The bottom panels consist of a slopegraph whereby each line represents a set of observations from an individual.

**Figure 7:**
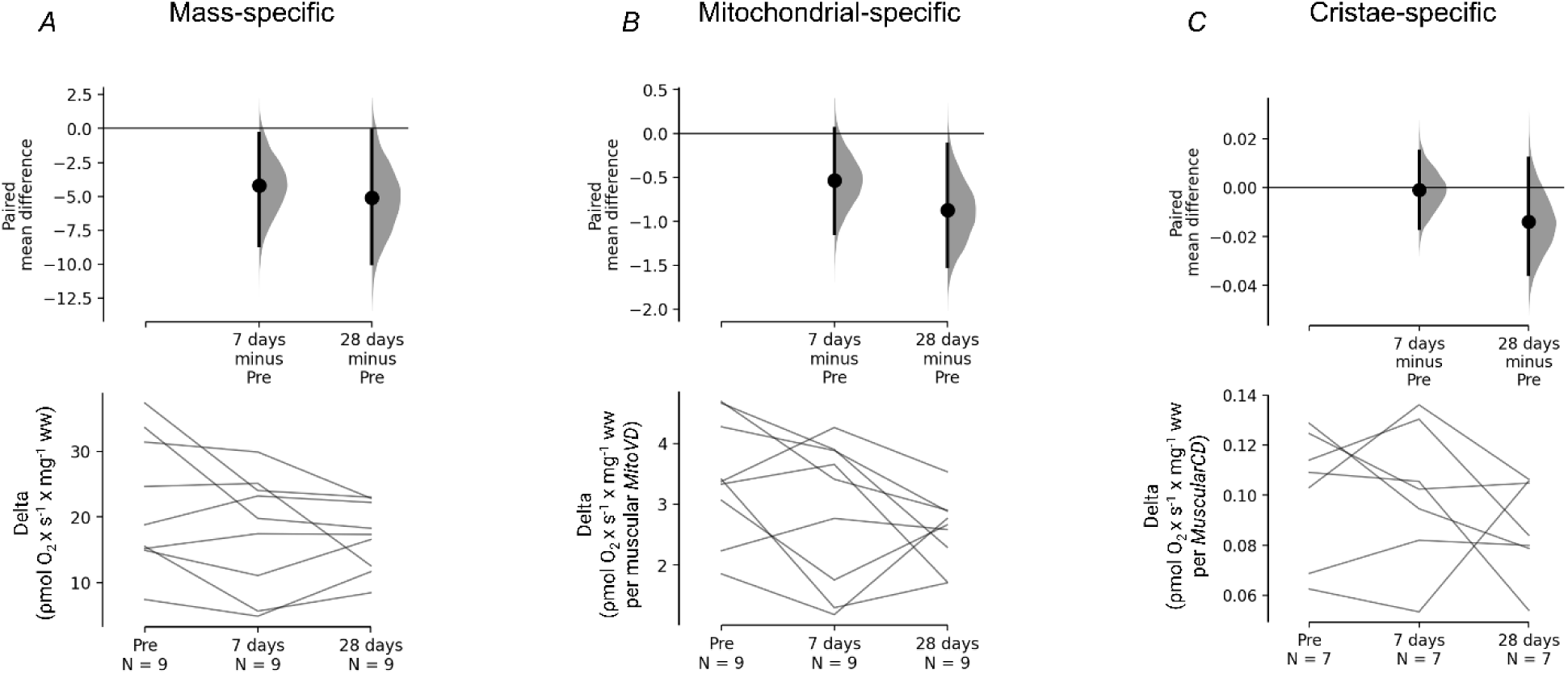
Effect of high altitude on mass-, mitochondrial- and cristae-specific mitochondrial coupling degree. *A,* mass-specific *FAO_ETF+CI* minus mass-specific *LEAK_ETF+CI* (delta); *B,* mitochondrial-specific *FAO_ETF+CI* minus mitochondrial-specific *LEAK_ETF+CI* (delta); *C,* cristae-specific *FAO_ETF+CI* minus cristae-specific *LEAK_ETF+CI* (delta). The paired mean difference between the timepoint before high-altitude exposure (Pre) and after high-altitude exposure (7 and 28 days) is displayed in a Cumming estimation plot. The top panels show the distribution of the bootstrapped-resampled paired mean difference between both the 7 days and 28 days timepoints and the Pre high-altitude timepoint. In each effect size half-violin plot, the black dot denotes the mean of the distribution, and the black vertical bars indicate 95% confidence intervals. The bottom panels consist of a slopegraph whereby each line represents a set of observations from an individual.

## Discussion

In this study we investigated how high-altitude exposure affects mitochondrial subcellular distribution, ultrastructure, respiratory control and intrinsic mitochondrial respiratory capacity by combining mass-specific mitochondrial respiration with *MuscularCD*. We found that exposure of sea-level residents to high altitude induced alterations in mitochondrial subcellular distribution, ultrastructure, and respiration, depending on the duration of exposure: Muscular *MitoVD* remained unchanged after 7 days at high altitude but increased following 28 days, which was driven by a rise in intermyofibrillar *MitoVD*. Despite this increase in muscular *MitoVD*, *MuscularCD* did not increase, and showed a slight reduction with acclimatisation due to a decrease in *MitoCD*. Although *MuscularCD* decreased, *OXPHOS_CII+CI+ETF*, when normalised to overall tissue sample mass, was slightly elevated. This remained unchanged when normalising mass-specific *OXPHOS_CII+CI+ETF* to muscular *MitoVD* but normalising to *MuscularCD* revealed a more pronounced increase in respiration. Despite being accompanied by a small increase in cristae-specific *Leak_CII+CI+ETF*, the difference between cristae-specific leak and maximal coupling respiratory states, indicated a significant higher degree of coupling between electron flow in the ETS and ATP production. Collectively, these results suggest an elevated intrinsic mitochondrial respiratory capacity after 7 days at high altitude, which persisted following 28 days.

### Effect of high altitude on the mitochondrial network and its subcellular distribution

As previously reported by Jacobs *et al*. (2016), muscular *MitoVD* increased following 28 days of high-altitude exposure (3454 m) in sea-level residents. In contrast, during a climbing expedition, no change in muscular *MitoVD* was observed after ∼13 days of trekking and 6 days at Everest Base Camp (5300 m) (Levett *et al*., 2012). Extending the exposure to ∼5-8 weeks, including brief ascents to higher elevations, such as attempts to summit Mt. Lhotse Shar (8398 m) or Mt. Everest (8848 m), resulted in a ∼20% decrease in muscular *MitoVD* two days after summit attempts (Levett *et al*., 2012) and 10-15 days upon return to sea level (Hoppeler *et al*., 1990). However, during expeditions confounding conditions are not easily controlled, and dietary availability is not necessarily the same as home and exercise patterns diverge from habitual activity. In an effort to address these confounding factors, MacDougall *et al*. (1991) used a live-in hypobaric chamber simulating an ascent to Mt. Everest over 40 days, where no change in muscular *MitoVD* was observed. Yet, despite access to plenty of varied palatable food and training equipment, caloric intake and daily exercise time were reduced during the confinement (Rose *et al*., 1988). In the present study, participants resided in the comfortable environment of the Jungfraujoch Research Station, where they were able to order groceries daily and replicated to the best of their ability normal sea level activities. Under these conditions, the typical body weight loss seen in climbing expeditions (Pugh, 1962; Boyer & Blume, 1984; Hoppeler *et al*., 1990; Wandrag *et al*., 2017) and hypobaric chamber confinement (Rose *et al*., 1988) was prevented, and, this less catabolic environment may have contributed to the observed ∼6% increase in muscular *MitoVD* in recreationally active sea-level residents.

When this effect size is considered in the context of mitochondrial plasticity observed in similar populations, the ∼6% increase in muscular *MitoVD* can be regarded as relatively modest. In comparison, a well-established stimulus such as exercise training elicits far greater increases in muscular *MitoVD*. For instance, previous studies have demonstrated a ∼40% increase following 6-8 weeks of training in untrained-recreationally active individuals (Hoppeler *et al*., 1985; Rösler *et al*., 1985; Montero *et al*., 2015), with increases as high as 116% when the training period is extended to 28 weeks (Kiessling *et al*., 1973). Despite the smaller change observed in this study, the ∼6% rise in muscular *MitoVD* may still represent a meaningful adaptation to the hypoxic environment, where tissue hypoxia necessitates an efficient mitochondrial network.

It has become increasingly clear that skeletal muscle mitochondria form a highly interconnected network that spans from the periphery to the interior of the muscle fibre (Bakeeva *et al*., 1978; Kirkwood *et al*., 1986; Glancy *et al*., 2015; Bleck *et al*., 2018). Considering a theory, which remains to be investigated experimentally, that suggests that the inner mitochondrial membrane may constitute a route of intracellular oxygen transport (Bakeeva *et al*., 1978; Skulachev, 1980; Pias, 2021) one important role of this interconnectivity can be to distribute oxygen to areas where it is needed within the muscle fibre. In this study, the observed increase in muscular *MitoVD* was driven by the intermyofibrillar pool of mitochondria. Since this pool surrounds the myofibrils (Eisenberg, 1983) such an expansion of the mitochondrial network could facilitate oxygen delivery to oxygen- demanding regions and may furthermore result in a faster distribution of oxygen by providing additional routes for oxygen transport i.e. in all, contributing to a more efficient mitochondrial network. Intermyofibrillar mitochondria has in fact been shown to exhibit a greater connectivity with the sarcoplasmic reticulum and lipid droplets than subsarcolemmal mitochondria in oxidative fibres via focused ion beam scanning electron microscopy (Parry *et al*., 2024). Using this technique in future studies would allow a spatial interpretation of the expansion of the intermyofibrillar pool of mitochondria induced by high-altitude exposure.

In contrast to the intermyofibrillar *MitoVD*, subsarcolemmal *MitoVD* decreased in response to high- altitude exposure. A reduction in the volume of the mitochondrial network within this space of the muscle fibre may serve to prevent excessive production of reactive oxygen species (ROS), which has been observed to increase during hypoxia (Guzy *et al*., 2005; Mansfield *et al*., 2005), although this remains a topic of debate. One possible explanation for the specific reduction in the subsarcolemmal *MitoVD* may relate to that the membrane potential can be transmitted along the mitochondrial network (Skulachev, 1971; Amchenkova *et al*., 1988), and the interconnectivity of the network thereby enables a distribution of potential energy from the edge all the way to the core of the skeletal muscle fibre (Skulachev, 1977; Bakeeva *et al*., 1978; Glancy *et al*., 2015; Parry *et al*., 2024). It is therefore speculated, that one role of the mitochondria, localised in the less spatially constrained subsarcolemmal region, is to serve as a reservoir of potential energy for the interior of the muscle fibre. In support, subsarcolemmal mitochondria, situated in large pools lateral to capillaries, demonstrates higher connectivity with neighboring mitochondria and showed signs of greater ETS activation compared to intermyofibrillar mitochondria (Parry *et al*., 2024). As a result, the subsarcolemmal pool may be reduced, prioritising the intermyofibrillar pool to ensure efficient energy distribution throughout the muscle fibre. However, given the wide 95% confidence interval associated with the effect size and the values included within the interval, this reduction in subsarcolemmal *MitoVD* is an uncertain finding that requires further investigations to confirm. Nevertheless, the intermyofibrillar pool of mitochondria seem to be given priority when the mitochondrial network adapts to high altitude.

### Effect of high altitude on mitochondrial network ultrastructure and intrinsic respiratory capacity

Following 28 days at high altitude, skeletal muscle *MitoCD* decreased by 5-10%, which contrasts with earlier assumptions that *MitoCD* remains constant (Hoppeler & Lindstedt, 1985; Hoppeler, 1986; Lindstedt *et al*., 1988; Weibel *et al*., 1991; Weibel & Hoppeler, 2005). In general, longitudinal studies on *MitoCD* are limited but several studies, investigating animal skeletal muscles, have suggested *MitoCD* plasticity based on micrograph observations (Gustafsson *et al*., 1965; Gollnick & King, 1969; Salmons *et al*., 1978). Later, when *MitoCD* was quantified, a change was found in rat skeletal muscles (Buser *et al*., 1982). Human cross-sectional studies also support *MitoCD* plasticity with higher values observed in endurance- (Nielsen *et al*., 2017; Schytz *et al*., 2024) and strength- trained athletes (Botella *et al*., 2023) compared to untrained individuals. Thus far, human longitudinal studies have failed to demonstrate an alteration in *MitoCD* (Lüthi *et al*., 1986; Nielsen *et al*., 2017). Thus, this is to the best of our knowledge the first longitudinal study to demonstrate changes in *MitoCD* in human skeletal muscle. The observed effect size in this study of 5-10% is substantial, knowing that untrained can differ from endurance athletes by ∼20% (Nielsen *et al*., 2017; Schytz *et al*., 2024) but it must be stressed that the associated 95% confidence interval include trivial effects meaning that high-altitude exposure may have only a minor impact on skeletal muscle *MitoCD*. Nevertheless, owing to the decreased *MitoCD*, the *MuscularCD* did not increase albeit the augmented muscular *MitoVD*. In fact, the effect size suggests a 5-10% decrease in *MuscularCD*, although the 95% confidence interval contain trivial effects.

A reduction in *MuscularCD* may help reduce the production of ROS. Mitochondrial ROS can be generated at specific sites within the ETS, such as Complexes I and III, where incomplete reduction of oxygen occurs (Zorov *et al*., 2014). Most ETS complexes are located in the cristae membrane (Gilkerson *et al*., 2003; Schlame, 2021) and, along with the ATP synthase complex, are thought to be involved in cristae formation (Mühleip *et al*., 2023). Although mitochondrial ROS has been shown to be important for stabilisation of hypoxia inducible factors (Mansfield *et al*., 2005), with Complex III playing a central role (Guzy *et al*., 2005), the observed decrease in *MuscularCD* during high- altitude acclimatisation may serve as a protective mechanism by reducing the amount of ETS complexes to avoid an imbalanced ROS production and prevent oxidative stress under hypoxic conditions.

The transfer of electrons through the ETS complexes to oxygen generates a proton gradient across the inner mitochondrial membrane, which drives conformational changes in the ATP synthase complex. This facilitates the binding of ADP and inorganic phosphate as well as the formation and release of ATP (Mitchell, 1966; Boyer, 1993; Saraste, 1999). Consequently, a reduction in *MuscularCD*, observed in this study, could potentially impair the ATP-producing capability of the mitochondrial network. However, mass- and mitochondrial-specific *OXPHOS_CII+CI+ETF* showed a slight elevation after high-altitude exposure, and when normalised to *MuscularCD*, a substantial increase was observed, suggesting intrinsic adaptations. This effect of high-altitude exposure may therefore have been masked in previous studies reporting only trivial effects on mass-specific maximal coupled respiration (Jacobs *et al*., 2013; Horscroft *et al*., 2017; Chicco *et al*., 2018). In contrast, Jacobs *et al*. (2012) found a ∼30% decrease in maximal coupled respiration normalised to CS activity. However, using the activity of a single protein as a biomarker for mitochondrial network changes is limited. Indeed, CS activity has been shown to be a poor biomarker of changes in muscular *MitoVD* during exercise interventions (Meinild Lundby *et al*., 2018).

Importantly, a greater cristae-specific *OXPHOS_CII+CI+ETF* does not necessarily equate to increased ATP production. To better assess the efficiency of oxygen consumption for ATP production, we subtracted cristae-specific *LEAK_CII+CI+ETF* from cristae-specific *OXPHOS_CII+CI+ETF*. This revealed a 30-40% higher mitochondrial coupling per *MuscularCD*, which was not evident in the mass- or mitochondrial-specific data. These findings suggests that high- altitude exposure enhances the intrinsic properties of the mitochondrial network. Additionally, since fatty acid oxidation requires more oxygen per ATP produced compared to carbohydrate oxidation (Krogh & Lindhard, 1920), the observed shift from fatty acid to carbohydrate metabolism after 28 days at high altitude indicates a more efficient use of available oxygen. However, while this shift improves oxygen efficiency at the skeletal muscle mitochondrial level, it does not occur during whole-body exercise at high altitude when performed at the same relative exercise intensity as at sea level (Lundby & Van Hall, 2002). Further research is needed to fully understand the broader implications of these high-altitude-induced adaptations.

### Concluding remarks

In this study, sea-level residents were exposed to 28 days of high altitude at the Jungfraujoch Research Station (3454 m), with skeletal muscle biopsies taken before ascent and after 7 and 28 days of exposure. Our findings demonstrated an altered subcellular distribution where an increase in the *MitoVD* of the markedly largest pool of mitochondria, the intermyofibrillar pool, led to a small increase in muscular *MitoVD*. This did however lead to a decrease in *MuscularCD* due to a substantial reduction in *MitoCD*. Despite a decrease in *MuscularCD*, mass-specific *OXPHOS_CII+CI+ETF* increased modestly after 28 days of high-altitude exposure. To assess whether this was attributed to intrinsic adaptations, *OXPHOS_CII+CI+ETF* was normalised to *MuscularCD*, which revealed a substantial increase in respiration following 7 days at high-altitude and remained elevated after 28 days. This was not evident when normalising *OXPHOS_CII+CI+ETF* to overall tissue sample mass or muscular *MitoVD*, highlighting that intrinsic adaptations may be overlooked in these measures. To examine intrinsic properties of the mitochondrial network, we evaluated the coupling efficiency between electron flow in the ETS and ATP production via oxygen kinetics by subtracting cristae-specific *Leak_CII+CI+ETF* from cristae-specific *OXPHOS_CII+CI+ETF*. This revealed a considerably greater degree of coupling after both 7 and 28 days of high-altitude exposure, suggesting improved mitochondrial respiratory efficiency. However, the 95% confidence intervals for the effect sizes did generally include trivial effects, indicating that further studies are required to substantiate these findings. If supported, our results suggest that high-altitude acclimatisation enhance mitochondrial network quality, favorable in an environment characterised by tissue hypoxia and maybe valuable knowledge to consider when planning training camps for athletes pursuing to enhance endurance performance.

## Acknowledgments

We express our sincere gratitude to the study participants for their willingness, time and effort devoted to this study.

## Author Contributions

CTS, CL, JN, NØ and RAJ contributed to the conception and design of the experiments. All authors contributed to the data collection, assembly, analyses and/or data interpretation. CTS and CL drafted the manuscript, while all authors edited and/or revised the manuscript. The final version of the manuscript was approved by all authors, and all agree to be accountable for all aspects of the work in ensuring that questions related to the accuracy or integrity of any part of the work are appropriately investigated and resolved. All persons stated as authors qualify for authorship, and all those who qualify for authorship are listed.

## Data availability

The data that support the findings of this study are available from the corresponding authors upon reasonable request.

